# Sex differences in crossover interference in house mice

**DOI:** 10.1101/2025.08.10.669551

**Authors:** Andrew P Morgan

**Affiliations:** Department of Medicine, University of North Carolina, Chapel Hill, NC 27514, USA

**Keywords:** meiosis, crossover interference

## Abstract

Meiotic recombination ensures the fidelity of chromosome segregation in most organisms with sexual reproduction. The distribution of crossovers along chromosomes is governed in part by interference, which prevents multiple crossovers from occurring in close proximity, though not all crossovers are subject to interference. Neither the factors that control the strength of interference, nor the extent to which they vary within and between species, are well understood. Here I confirm that crossover interference is stronger in male than in female meiosis in house mice (*Mus musculus*), provide the first estimate of the proportion of non-interfering crossovers in female mice, and show that this proportion is lower than in males. Interference is stronger on shorter chromosomes in both sexes, but the frequency of non-interfering crossovers is similar across the range of chromosome size. Together with evidence that interference varies across strains and subspecies, my results provide a foundation for studying the evolution and sexual dimorphism in this important feature of meiosis in mice.

## Introduction

Meiotic recombination is essential for faithful chromosome segregation in most sexually-reproducing organisms (Hassold and Hunt 2001). Both the number and the spatial distribution of crossovers are tightly regulated, but vary within and between species. At fine scale, the position of crossovers is determined in part by the location of double-strand breaks, which in turn are associated with specific sequence motifs and/or epigenetic marks which differ by taxa (reviewed in Paigen and Petkov (2010)). At the chromosome scale, crossovers tend to be spaced more evenly than expected by chance. This observation dates back to the first linkage maps inferred from *Drosophila* (Sturtevant 1913, 1915) and is known as crossover interference.

The mechanisms and evolutionary significance of crossover interference remain incompletely understood (Hunter 2015). Muller (1916) first speculated that even spacing of crossovers might promote orderly chromosome segregation. There is now ample evidence that aberrant placement of crossovers with respect to the centromere, telomere or to each other contributes to nondisjunction (Hassold *et al.* 1991; Koehler *et al.* 1996; Hassold and Hunt 2001). Others have proposed that interference arises as a consequence of processes that ensure at least one crossover per bivalent (Jones 1984) and maintain the total number of crossovers per meiosis within some optimum range (Martini *et al.* 2006; Cole *et al.* 2012). Further, a subset of crossovers are also processed via a distinct pathway that is not subject to interference (Santos *et al.* 2003; Higgins *et al.* 2004; Guillon *et al.* 2005; Holloway *et al.* 2008). Like other features of the recombination landscape (Morelli and Cohen 2005; Sardell and Kirkpatrick 2020), both the magnitude of interference and the frequency of non-interfering crossovers vary between sexes in several mammal species (Otto and Payseur 2019). Comparative studies are hindered by the relative paucity of empirical estimates of interference parameters in both sexes. Recombination tends to be much easier to study in the male than in the female germline – spermatocytes are usually more readily obtained than oocytes, and breeding many progeny from one male is often less resource-intensive than breeding the same number of progeny from one female. Our knowledge of meiosis in mammals is thus biased towards the male germline.

Here I use a large panel of 8-way intercross pedigrees from the Collaborative Cross (CC) (Churchill *et al*. 2004) to provide sex-specific estimates of the strength of crossover interference and the proportion of non-interfering crossovers in both sexes of house mice (*Mus musculus*). The CC is an excellent vehicle for studying recombination (Liu *et al.* 2014) because it comprises hundreds of informative meioses through both sexes in a randomized genetic background with balanced contributions from 8 founder strains representing all the major subspecies of mice (*M. m. domesticus, M. m. musculus* and *M. m. castaneus*). I confirm that interference is stronger in males than females and show for the first time that the proportion of non-interfering crossovers is also greater in males than females. Longer chromosomes show stronger interference than shorter chromosomes in both sexes, but no difference in proportion of non-interfering crossovers.

## Results and discussion

Fully-phased autosomal haplotypes were obtained from one male and one female offspring from each of 237 intercross families with the structure shown in **Figure 1A**. Briefly, the 8 founder strains (G_0_ generation, not shown) were randomly intercrossed to create the G_1_ hybrids, which were again intercrossed to create the G_2_ hybrids. The offspring of the G_2_ pairs are denoted the G_2_:F_1_ and their genomes are a readout of the meioses in the G_1_ and G_2_ animals. Pedigree constraints allow each haplotype junction to be assigned to a crossover event in exactly one of 8 meioses, so the 474 offspring provide at least partial information for the outcome of 1 896 meioses (see Liu *et al.* (2014)). (Crossovers in the G_1_ meioses are only observed if they are transmitted to the G_2_:F_1_.) I limit my attention to the 948 fully observed meioses in the G_2_ generation, which resulted in a total of 12 191 crossovers (6 537 in female and 5 654 in male meioses). Of 18 012 transmitted chromosomes, 7 457 (41.4%) were non-recombinant, 8 950 (49.7%) had exactly one crossover and 1 605 (8.9%) had two or more crossovers. An average of 13.8 crossovers were transmitted per meiosis in females and 11.9 in males. On chromosomes with more than one crossover, the mean distance between adjacent crossovers was 44.8 cM (SD 14.2 cM) in females and 44.4 cM (SD 13.0 cM) in maless, on the sex-specific genetic maps estimated from this population (**Figure 1B**).

**Figure 1.**
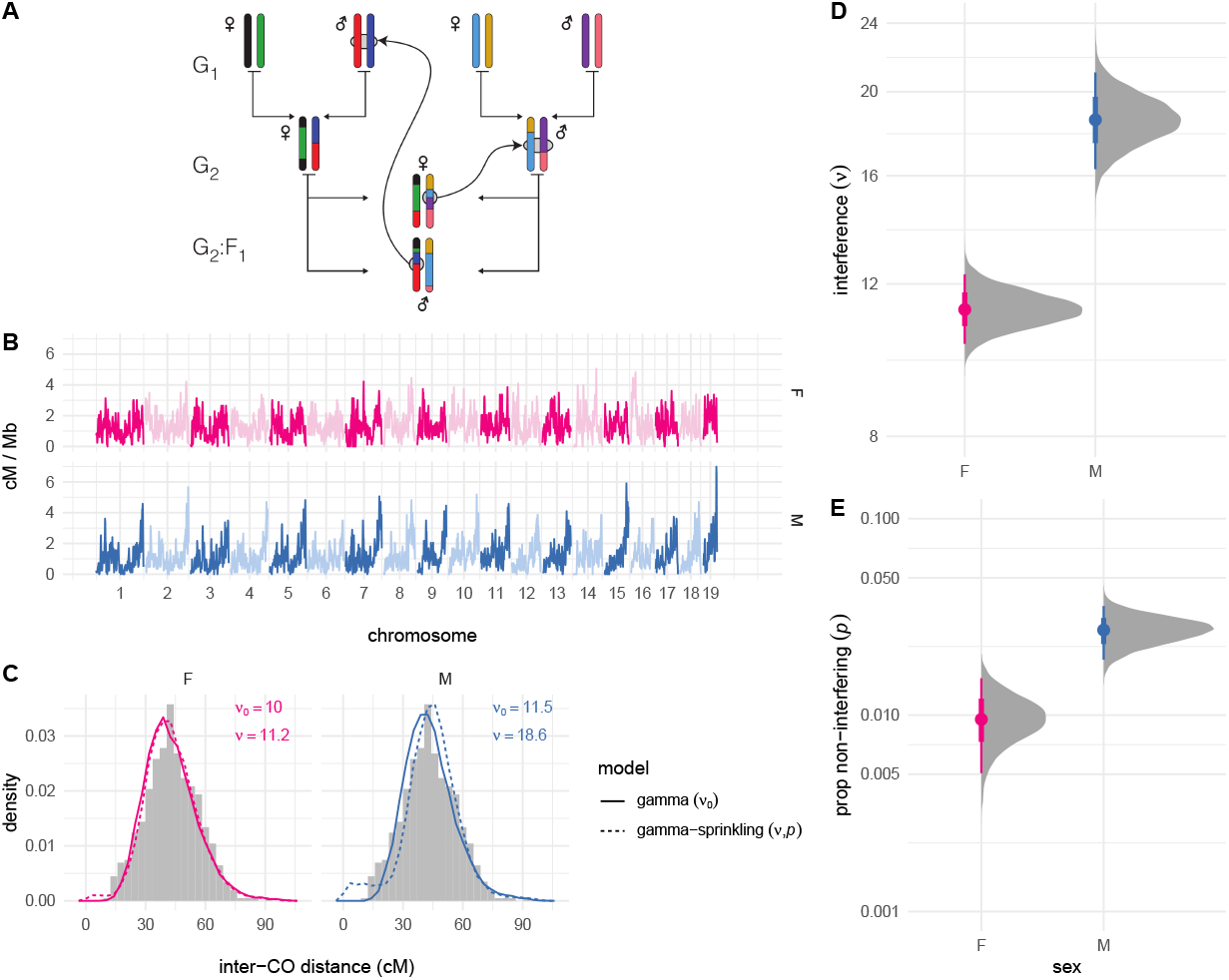
Estimating crossover interference genome-wide. (**A**) Pedigree of a representative intercross family. Founder haplotypes are color-coded, and each family is uniquely defined by the order of founder strains in the first generation. Crossovers identified in the G_2_:F_1_ offspring can be assigned to exactly one meiosis. (**B**) Sex-specific genetic maps, smoothed over 1 Mb windows. (**B**) Empirical distribution of inter-crossover distances (histograms) compared to simulations (lines) using posterior mean values for *ν*_0_ (gamma model) or {*ν, p*} (gamma-sprinkling model). (**D**,**E**) Posterior distribution of global sex-specific estimates of *ν* and *p*. Solid dots show posterior mean; bars show 25% − 75%ile (thick) and 2.5% − 97.5%ile (thin) quantile intervals.

Sex-specific interference parameters were estimated under the gamma (Broman and Weber 2000) and “gamma-sprinkling” (Housworth and Stahl 2003) models of crossover interference using an extension of a Bayesian procedure I have described previously (Morgan and Payseur 2024) (**Figure 1C-E**). The gamma model quantifies strength of interference via a single parameter (here denoted *ν*_0_). The gamma-sprinkling model treats crossovers as a mixture of two processes: with interference, under the gamma model (with probability 1 − *p*); or without interference, equivalent to the gamma model with *ν* ≡ 1 (with probability *p*). Importantly, these are statistical abstractions and not mechanistic models. See **Supplementary note 1** for motivation and details. Under the simpler gamma model, interference was stronger in males (*ν*_0,male_ = 11.5 [95% HPDI 10.5 − 12.5]) than in females (*ν*_0,female_ = 10.0 [95% HPDI 9.3 − 10.8]). At the maximum-likelihood estimate, the gamma-sprinkling model provided better fit to the data (−2 * log-likelihood ratio = 84.2, *p* = 5.2 *×* 10^−19^, likelihood-ratio test; ΔAIC = 80.2). Subsequent results are from the the model including non-interfering crossovers since it is both biologically plausible and statistically supported. Interference was 1.6 times stronger in males (*ν*_male_ = 18.6 [95% HPDI 16.3 − 21.1]) than in females (*ν*_female_ = 11.2 [95% HPDI 10.2 − 12.3]). The proportion of crossovers that escape interference was also 2.9 times greater in males (*p*_male_ = 0.027 [95% HPDI 0.019 − 0.036]) than in females (*p*_female_ = 0.0094 [95% HPDI 0.0054 − 0.016]). As a posterior predictive check, the distribution of inter-crossover distances from simulations under the gamma and gamma-sprinkling models was qualitatively similar to the observed distribution (**Figure 1C**).

Many prior studies have shown that crossover interference is heterogeneous across chromosomes. I extended my model to allow chromosome-specific values of *ν* and *p* but was not able to obtain stable estimates due to limited sample size. Results from a reduced model without non-interfering crossovers were more stable and showed strong negative correlation between *ν* and chromosome length in both sexes (Spearman’s *ρ* = −0.39, *p* = 0.017; **Figure S1**). I reasoned that I could at least partially capture this signal by aggregating over chromosomes of similar size, and therefore divided them into three groups, each comprising about one-third of the cumulative genetic map. Results are shown in **Figure 2**. Interference was clearly stronger on shorter relative to longer chromosomes – 1.5 times (95% HPDI 1.1 − 1.9) stronger in females and 2.4 times (95% HPDI 1.6 − 3.7) stronger in males. The proportion of non-interfering crossovers was similar across the range of chromosome size (just 0.96 times greater in females and 1.0 times greater in males, not significantly different from unity). Sex differences in interference parameters were preserved across the range of chromosome size, though the difference in strength of interference between short and long chromosomes was greater in males. Considering female meioses only, interference on the X chromosome was similar in magnitude to that on autosomes of similar size (**Figure S2**).

**Figure 2.**
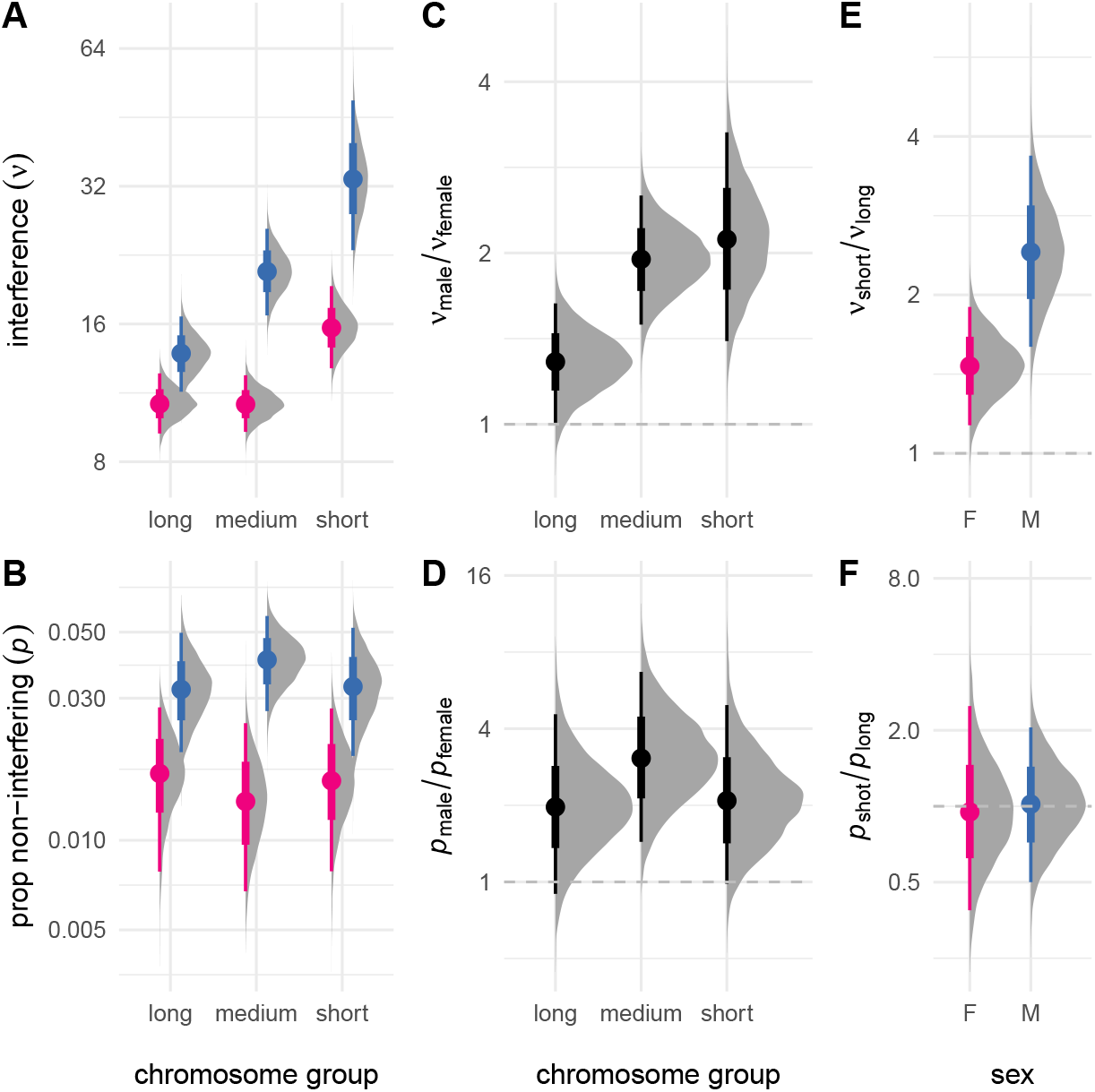
Interchromosomal variation in crossover interference. (**A**) Strength of interference estimated for chromosomes binned by size (“long”, chromosomes 1-5; “medium”, chromosomes 6-12; “short”, chromosomes 13-19). (**B**,**C**) Proportion of non-interfering crossovers. (**C**) Sex contrasts for *ν* and *p*, expressed as ratios. (**D**,**E**) Sex-specific contrasts between *ν* and *p* for “short” and “long” chromosomes.

Previous studies in mice based on linkage mapping of chromosome 1 (Petkov *et al.* 2007) and cytological analyses (de Boer *et al.* 2006; Peterson and Payseur 2021) have shown stronger interference in males than females. I confirm those findings. my estimate of *ν*_female_ = 11.2 was nearly identical to the only previous estimate from linkage data (11.3) in (*Mus musculus × Mus spretus*) F_1_ females (Broman *et al.* 2002). As with cattle (Wang *et al.* 2016), pigs (Brekke *et al.* 2025), domestic dogs (Campbell *et al.* 2016) and humans (Campbell *et al.* 2015), I observed an inverse relationship between (genetic) chromosome size and chromosome-level interference in both sexes. This results in a negative correlation between the genome-wide strength of interference and the average genetic length of a chromosome arm across species (Spearman’s *ρ* = −0.66, *p* = 0.022; **Figure S3**), though there is no such relationship with the proportion of non-interfering crossovers (Spearman’s *ρ* = 0.28, *p* = 0.38). The proportion of non-interfering crossovers in mice was about half as large as in these other species (**Table 1**). At least in this small group of species, there is no consistent pattern as to whether interference is stronger in males or females; in the sex with the longer or shorter genetic map; or in the sex with more or fewer non-interfering crossovers.

**Table 1:** Sex-specific interference parameters for available mammal species. cM, length of the autosomal recombination map. N, haploid chromosome number. FN, number of haploid chromosome arms (“fundamental number”, (Matthey 1945)).

| species | sex | cM | N | FN | $\nu$ | p | reference |
| --- | --- | --- | --- | --- | --- | --- | --- |
| mouse | M | 1221 | 19 | 19 | 18.60 | 0.0268 | this study |
|  | F | 1355 |  |  | 11.20 | 0.00937 |  |
| dog | M | 1816 | 38 | 38 | 14.05 | 0.055 | Campbell *et al.* (2016) |
|  | F | 2162 |  |  | 30.64 | 0.035 |  |
| cow (Holstein) | M | 2519 | 29 | 58 | 8.70 | 0.0834 | Wang *et al.* (2016) |
|  | F | 2372 |  |  | 9.96 | 0.1002 |  |
| cow (Jersey) | M | 2369 | 29 | 58 | 6.70 | 0.0220 | Wang *et al.* (2016) |
|  | F | 2229 |  |  | 7.55 | 0.0289 |  |
| pig | M | 1915 | 18 | 32 | 8.54 | 0.049 | Brekke *et al.* (2025) |
|  | F | 2664 |  |  | 6.45 | 0.043 |  |
| human | M | 2708 | 23 | 46 | 8.93 | 0.067 | Campbell *et al.* (2015) |
|  | F | 4355 |  |  | 7.19 | 0.078 |  |

Why does crossover interference differ between sexes? One possibility is that the physical process(es) that constitute interference, whatever they are, scale with the length of the synaptonemal complex (SC). The SC is longer in females in humans (Tease and Hultén 2004) and mice (de Boer *et al.* 2006), and crossover counts are positively correlated with SC length within and across individuals in many species (Lynn *et al.* 2002; de Boer *et al.* 2006; Dumont and Payseur 2011; Wang *et al.* 2019). Petkov *et al.* (2007) showed that the coefficient of co-incidence, a measure of interference between pairs of loci, depends on distance between loci on the SC (in µm) rather than in genomic coordinates (base pairs) in mice. This predicts that interference will be stronger in the sex with shorter SC relative to genome size and shorter genetic map. Humans, mice and pigs fit this pattern; dogs and cows do not. The relationship between SC length and recombination rate is very well-conserved while the direction of sex differences in interference is not. Factors beyond SC length must therefore be at play and it seems likely that some of these are lineage-specific. Since crossovers tend to be suppressed near centromeres (Hunter 2015) but the number of crossovers per meiosis is proportional to the number of chromosome arms (Pardo-Manuel de Villena and Sapienza 2001), lineage-specific features of chromosome organization and karyotype may contribute. So too may differences in life history: longer generation time means longer duration of meiotic arrest, and may favor weaker interference in females for optimal positioning of crossovers for secure attachment to the spindle (Kong *et al.* 2004; Campbell *et al.* 2015). It is tempting to speculate that sex differences in crossover interference are related to the unique susceptibility of the female germline to meiotic drive – to the extent that weaker interference alters the probability of a crossover between the centromere and a driving locus, it could oppose the spread of the driver if it acts at meiosis I, or favor spread if it acts at meiosis II (Brandvain and Coop 2012). The direction and magnitude of effect would depend on the genetic position, mechanism of action, and fitness effect of the driving allele. Finally, sex differences in interference may arise indirectly from the balance between sex-specific (and possibly sexually-antagonistic) selective pressures on recombination rate (Mank 2009; Dumont 2017) and mechanisms that ensure crossover homeostasis.

What explains the divergent relationship between *ν* and *p* on chromosome size? The biophysical constraints on the maximum number of crossovers and minimum spacing of crossovers compatible with successful chromosome segregation, may be such that chromosomes below some threshold size can accommodate one and only one crossover. That corresponds to complete interference (*ν* → ∞), and indeed, small chromosomes tend to have larger values of *ν* across species. If the non-interfering pathway is not subject to the same constraints, I predict that *p* would be uncorrelated with chromosome size.

Some caveats do apply to these results. First, by using haplotypes transmitted in live offspring as the substrate for analysis, I limit my study to viable gametes only. To the extent that interference is systematically different in gametes which are eliminated by meiotic checkpoints or cannot produce a viable zygote, my findings may be biased. Second, it is not clear how to connect the treatment of interference as a statistical phenomenon to an underlying molecular process. Mechanistic models such as the beam-film model (Kleckner *et al*. 2004; Zhang *et al.* 2014) offer an alternative perspective. In this model, the chromosome axis is treated as a thin “film” on the surface of a flexible “beam” under some tensile stress, across which are arrayed precursor sites. A “crack” (corresponding to a crossover) can form at a precursor site, which relieves tensile stress locally and inhibits further cracks (crossovers) nearby. The effect decays over space, so at sufficient distance away from the first crack, another can form, repeating the process. This continues until tensile stress has been dissipated to the extent that no more cracks are induced. Several of the key features of crossover patterning – interference, obligate crossover, crossover homeostasis and the genetic map itself – arise naturally from this model. Zhang *et al.* (2014) show that the distribution of inter-crossover distance is sensitive to the number of precursor sites (double-strand breaks), the probability that a precursor site will mature into a crossover and the distance over which tensile stress decays. The latter of these is probably most similar to the statistical description of interference. Simulation results from with the beam-film model imply that the quantity *ν* is actually a function of multiple distinct layers of regulation. Fitting the beam-film model to empirical data is not straightforward and, importantly, requires knowing the position of all crossovers on a chromosome (as by cytological measurements on gametocytes) and not just the random subset that are transmitted to progeny. The gamma and gamma-sprinkling models remain useful for describing interference from pedigree data in a common frame of reference across different organisms.

Finally, I note that the while *p* is an estimate of the of the proportion of non-interfering crossovers in aggregate, some independent validation is needed. These “class II” crossovers are resolved by distinct enzyme complexes (reviewed in Gray and Cohen (2016)) but experimental methods to identify them are less mature than those for the class I crossovers that predominate in most eukaryotes studied. The gamma-sprinkling model generally has better fit to empirical data than simpler the gamma model, as shown in this manuscript and elsewhere (Otto and Payseur 2019), but it is possible that the parameter *p* just adds some flexibility to the lower tail of the distribution of inter-crossover distances rather than directly estimating the proportion of non-interfering crossovers.

## Materials and methods

This work re-analyzes published crossover data from the mouse Collaborative Cross (CC) project (Churchill *et al.* 2004). Briefly, the CC is a panel of recombinant inbred mouse lines (RILs), each of which was initiated with an 8-way intercross followed by inbreeding. One male and one female of the last outbreeding generation in a subset of 237 lines were genotyped using a high-density SNP array and used to characterize the genome of the incipient RILs. Mouse breeding, DNA extraction, genotyping, haplotype inference, and construction of the recombination map are described in detail in Liu *et al.* (2014). Coordinates of haplotype segments were defined on the sex-specific genetic map estimated from this cross.

Crossover interference was analyzed under the “gamma-sprinkling” (or Housworth-Stahl) model. Some proportion *p* ∈ (0, 1) of crossovers are modeled as non-interfering, while the remaining 1 − *p* are subject to interference under the gamma model (Broman and Weber 2000) in which the strength of interference is represented by the single unitless parameter *ν >* 0. (Note that my parameterization assumes that *p* is strictly positive, though it can be arbitrarily small. This differs from some other implementations.) I allow both *ν* and *p* to vary across groups of meioses (indexed by *i* = 1, … *m*) and groups of chromosomes (indexed by *j* = 1, … *n*), using an extension of the hiearchical Bayesian approach described in Morgan and Payseur (2024). Let group-specific parameters *ν*_*ij*_, *p*_*ij*_ be defined as follows:

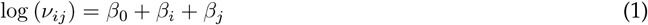

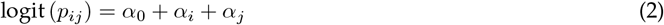

where *β*_0_ is a global intercept term (to apply some shrinkage) and the *β*_*i*_ and *β*_*j*_ are subject- and chromosome-specific effects. The prior distribution is given by 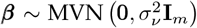. That is, group-specific *β*_*i*_ are assumed uncorrelated. The *α* terms are defined similarly. The likelihood of a set of transmitted chromosome segments, given the parameter vector ***θ*** = (***β, α***), was calculated with the `C` routine `R_stahl_loglik` from the `R` package `xoi` v0.72 (Broman 2023). Inference was done by a simple Metropolis-coupled Markov chain Monte Carlo (MCMC) sampler, with proposal distribution tuned to achieve a rejection rate of around 0.4. The sampler was run for 25 000 iterations, of which the first 5 000 were discarded as burn-in. Quantities of interest were calculated directly from the posterior distributions, with summary statistics calculated using `R` package `coda` v0.19-4.1.

## Data availability

Processed data underlying key analyses in this manuscript will be available on Figshare:

`doi:10.6084/m9.figshare.29876591`

Analysis code is available on Github:

https://github.com/andrewparkermorgan/mouse_crossover_interference_2

## Acknowledgements

I thank Bret Payseur for helpful discussions and the anonymous reviewers for constructive suggestions on sharpening the manuscript.

## Funding

Support was provided by the following grants from the National Institutes of Health: P50 GM076468 (Gary A Churchill) and F30 MH103925 (Andrew Morgan).

## Conflicts of interest

None of the authors have any competing financial interests in the work described in this manuscript.

## Supplementary material

**Supplementary note 1**. Description of statistical models for crossover interference.

**File S1**. Table of 21 993 crossovers (COs) reported in (Liu *et al.* 2014). Columns are as follows.

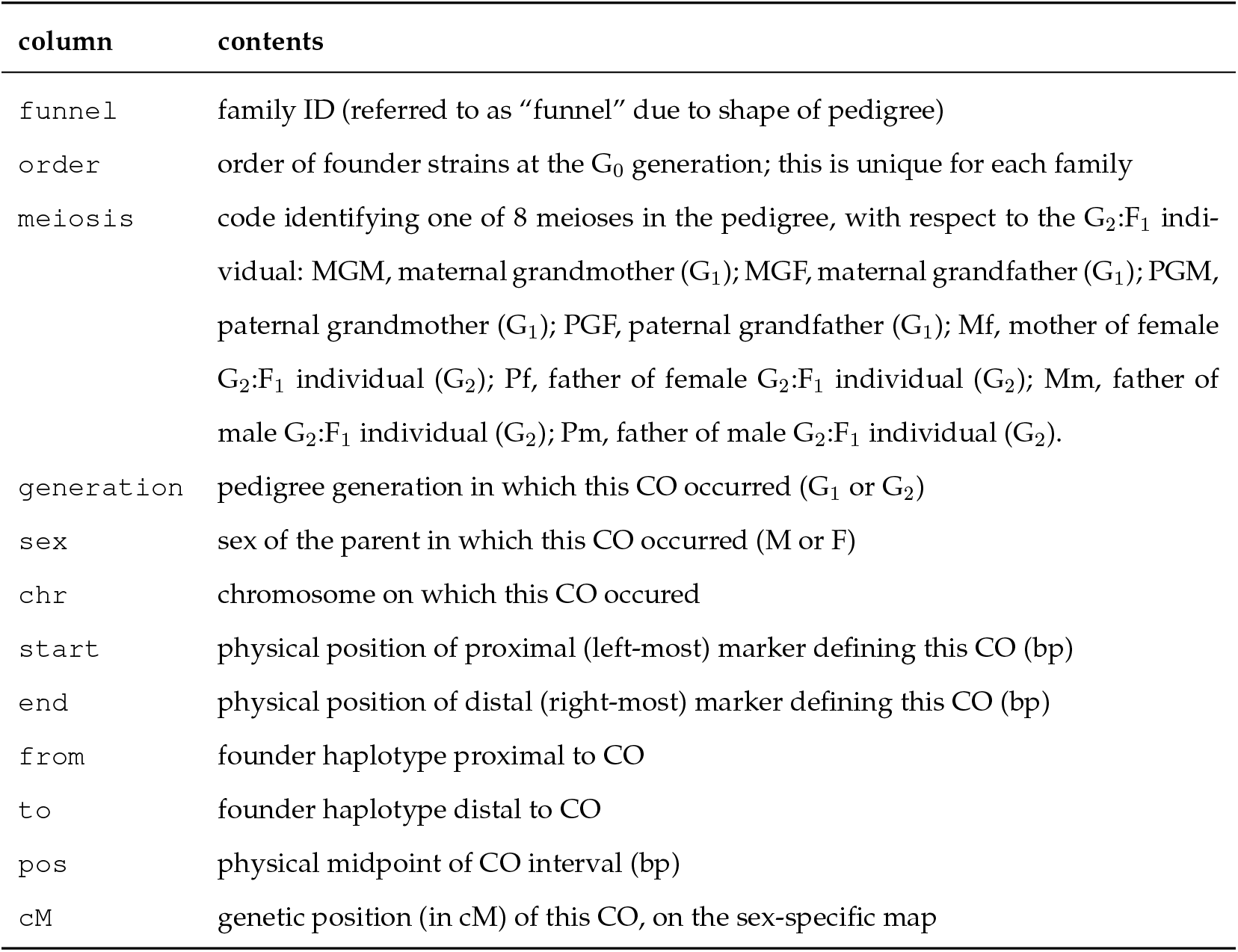

| column | contents |
| --- | --- |
| funnel | family ID (referred to as “funnel” due to shape of pedigree) |
| order | order of founder strains at the $G_0$ generation; this is unique for each family |
| meiosis | code identifying one of 8 meioses in the pedigree, with respect to the $G_2:F_1$ individual: MGM, maternal grandmother ($G_1$); MGF, maternal grandfather ($G_1$); PGM, paternal grandmother ($G_1$); PGF, paternal grandfather ($G_1$); Mf, mother of female $G_2:F_1$ individual ($G_2$); Pf, father of female $G_2:F_1$ individual ($G_2$); Mm, father of male $G_2:F_1$ individual ($G_2$); Pm, father of male $G_2:F_1$ individual ($G_2$). |
| generation | pedigree generation in which this CO occurred ($G_1$ or $G_2$) |
| sex | sex of the parent in which this CO occurred (M or F) |
| chr | chromosome on which this CO occurred |
| start | physical position of proximal (left-most) marker defining this CO (bp) |
| end | physical position of distal (right-most) marker defining this CO (bp) |
| from | founder haplotype proximal to CO |
| to | founder haplotype distal to CO |
| pos | physical midpoint of CO interval (bp) |
| cM | genetic position (in cM) of this CO, on the sex-specific map |

## Additional notes

Physical positions are reported on the mm9/GRCm37 assembly, as this was used during the original analysis, without liftover to the current mm39 assembly. Only genetic positions are actually used in this manuscript. Founder strains are reported as 1-letter codes: A = A/J; B = C57BL/6J; C = 129S1/SvImJ; D = NOD/ShiLtJ; E = NZO/HlLtJ; F = CAST/EiJ; G = PWK/PhJ; H = WSB/EiJ.

**Figure S1:**
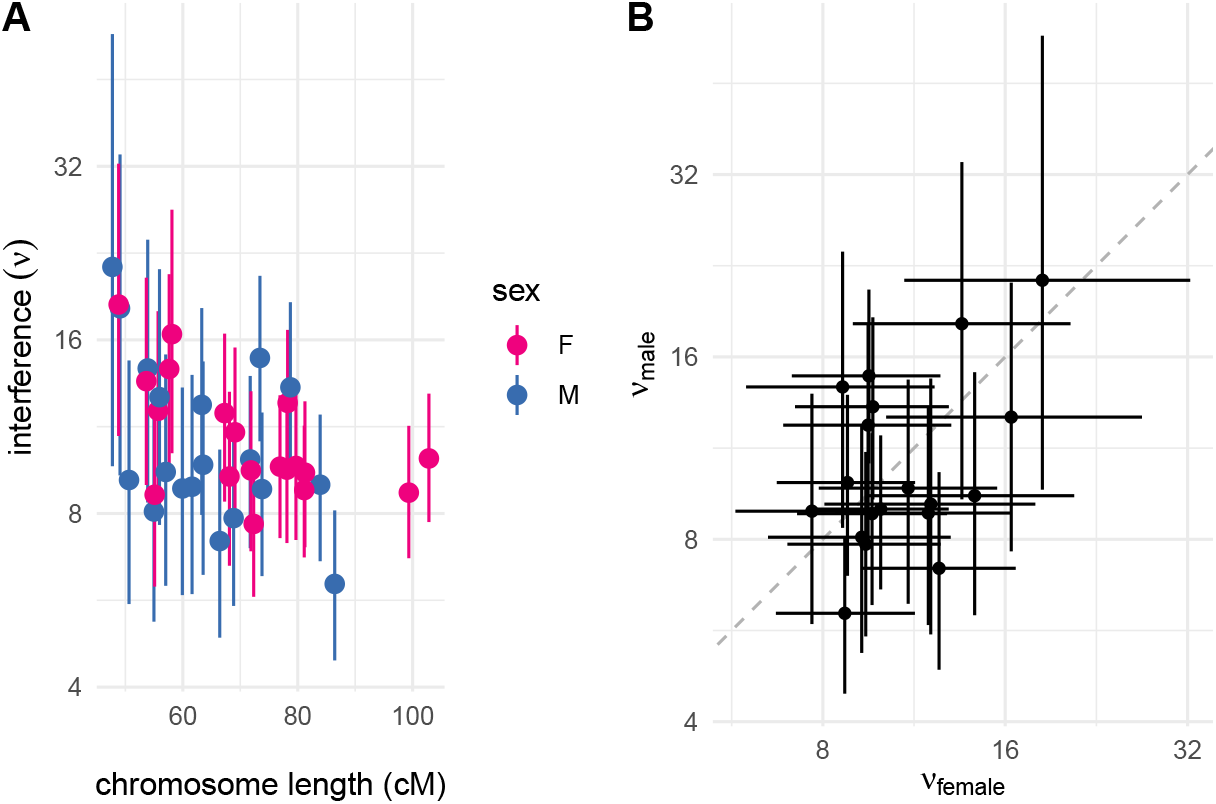
Chromosome-specific interference parameters under the simple gamma model without non-interfering crossovers. (**A**) Chromosome-specific values of *ν* with 95% HPDIs plotted against genetic chromosome length. (**B**) Comparison of male versus female estimates of *ν* for each chromosome. Dotted reference line passes through the intercept and has slope 1.

**Figure S2:**
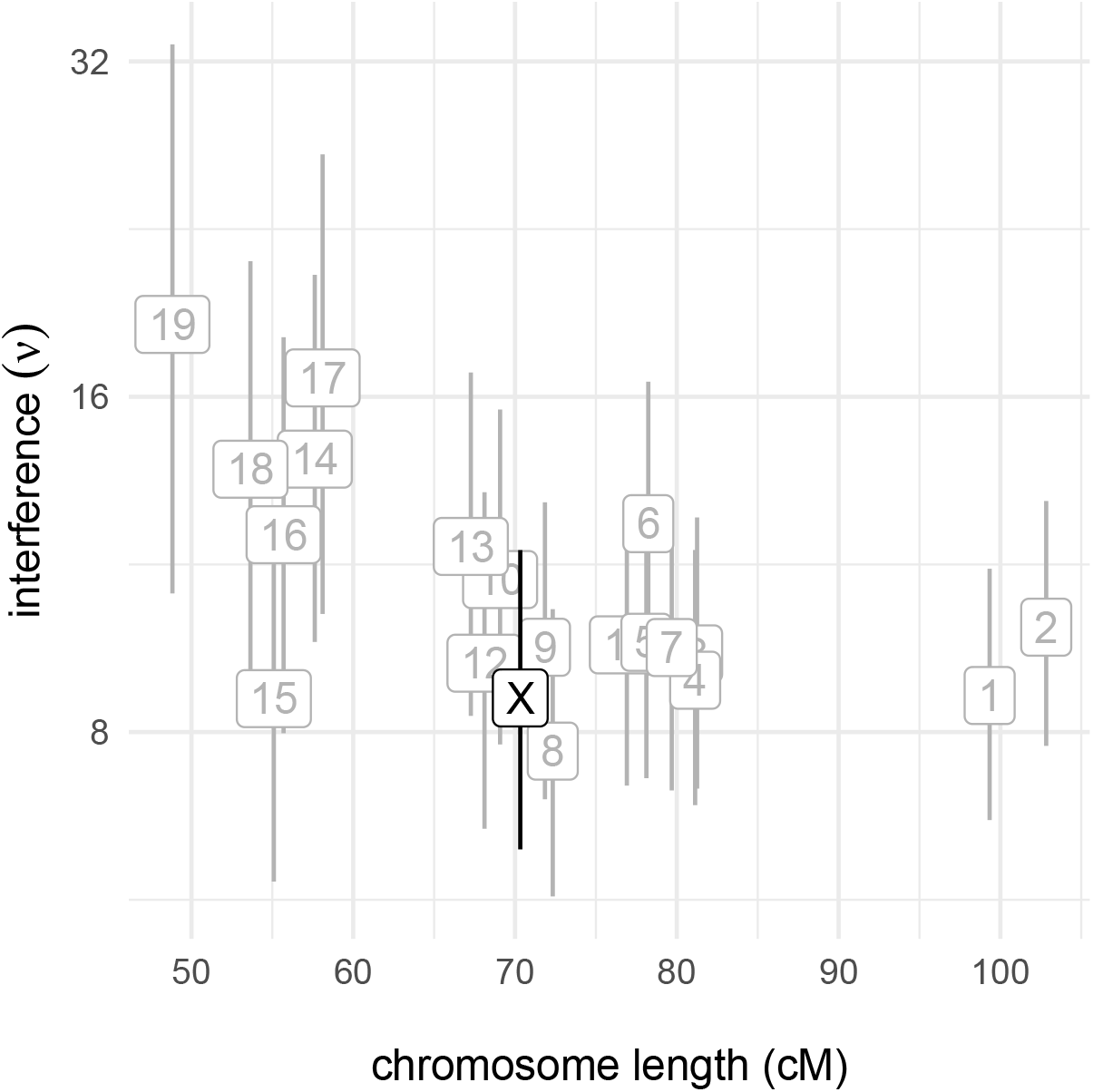
Chromosome-specific interference parameters under the simple gamma model without non-interfering crossovers, in females only. Chromosome-specific values of *ν* with 95% HPDIs plotted against genetic chromosome length, for labelled autosomes (grey) and the X chromosome (black).

**Figure S3:**
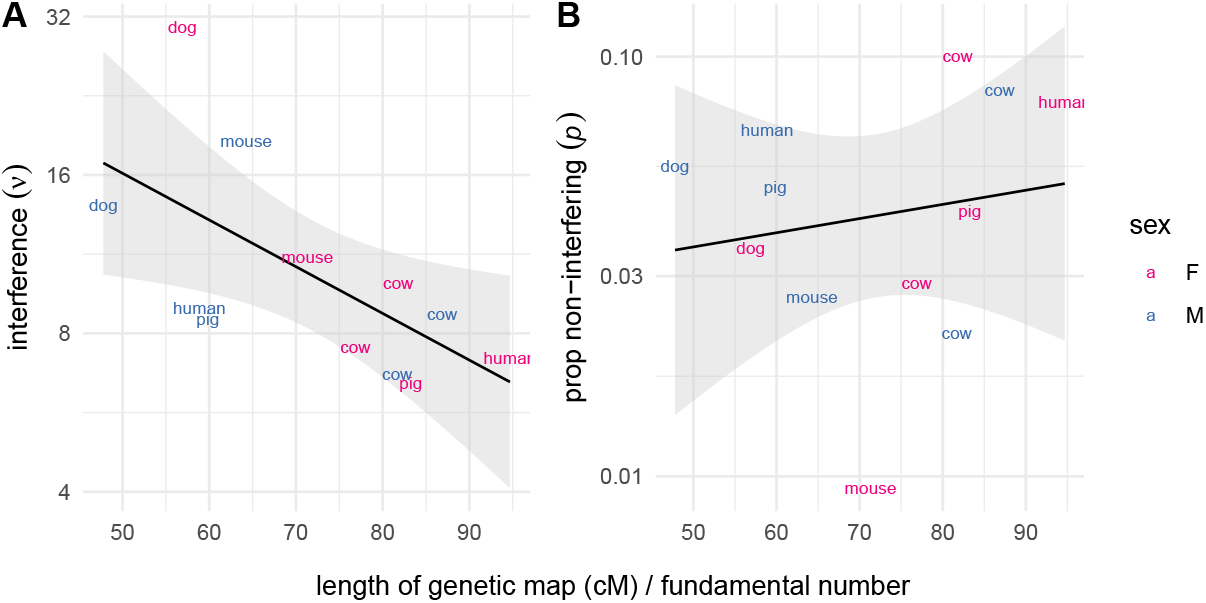
Interference vs length of genetic map. Sex-specific values of *ν* (**A**) and *p* (**B**) plotted against the average genetic length of a chromosome arm in 5 mammal species.

## Notes

http://dx.doi.org/10.6084/m9.figshare.29876591

https://github.com/andrewparkermorgan/mouse_crossover_interference_2

### Competing Interest Statement

The authors have declared no competing interest.

### Summary of Updates

(1) Addition of supplementary note detailing the statistical model for crossover interference (note: this is review of prior literature, not new intellectual contribution) (2) Addition of results from gamma model as well as gamma-sprinkling model (3) Discussion of gamma model in comparison to the (mechanistic) beam-film model for interference

