## Supplementary note for "Sex differences in crossover interference in house mice"

Andrew P Morgan

October 20, 2025

In this note I review the motivation for the gamma and gamma-sprinlking (aka gamma-escape, Housworth-Stahl or two-pathway) models of crossover interference, and how they are estimated from genetic data. Readers should refer to the original literature (Broman and Weber 2000; Housworth and Stahl 2003) for full mathematical detail. It should be emphasized that these are *statistical abstractions* whose purpose is to fit the observed distribution of the lengths of transmitted haplotype segments. Although the models account for some very biological features of meiosis – the linear organization of chromosomes and segregation of homologues – they are otherwise agnostic to the actual biophysical mechanisms that generate the phenomenon we call interference.

#### Gamma model

The gamma model is motivated by the fact that observed crossovers are downstream of the actual object of interference, namely the positions of chiasmata. Chiasmata are treated as arising via a stationary renewal process along an infinintely-long chromosome, for which the waiting time between events follows a gamma distribution with shape $\nu$ and rate $2\nu$ (see **Figure A**). Since each chromatid is transmitted with probability $\frac{1}{2}$, the chiasmata are "thinned" to produce the observed crossovers. The distances between chiasmata (denoted $x_i$) correspond to the waiting time of the renewal process, and follow a gamma distribution (denoted $\Gamma(\cdot)$). (Since chromosomes are not infinitely long, there is a further complication in handling the length of segments between the chromosome ends and the first and last chiasmata, the details of which are not shown here.) The observed inter-crossover distances (denoted $y_i$), on the genetic and not the physical map, are produced by the thinning process, and their distribution is a convolution of gamma distributions.

To estimate $\nu$ from genetic data, the meiotic products are converted to a collection of haplotype segments of four types, depending on whether the start- and end-points are either the end of a chromosome or a crossover.

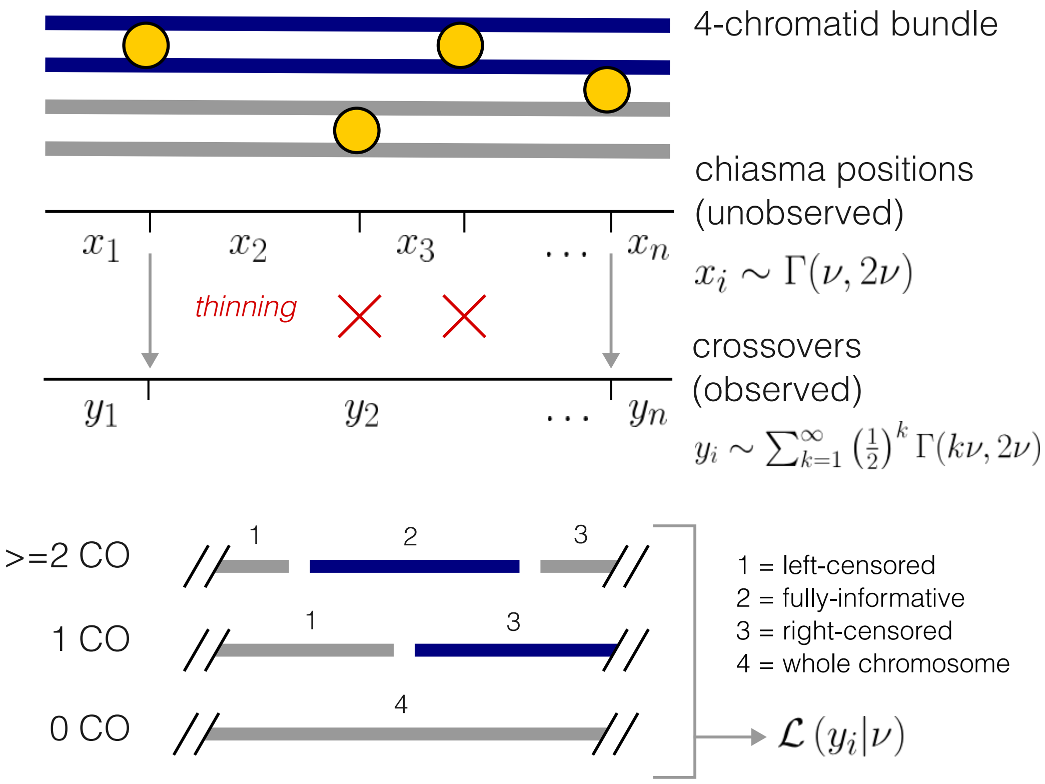


Figure A: **Overview of the gamma model of crossover interference.**

Segments of type 1 are bounded on the left by the proximal end of a chromosome, and on the left by a crossover; type 2, flanked by crossovers; type 3, bounded on the left by a crossover and on the right by the end of a chromosome; and type 4, whole non-recombinant chromosomes. Segments of types 1, 2 and 4 are treated as censored on the left, right or both ends, respectively – were the chromosome longer, another crossover eventually would have occurred. Due to stationarity, types 1 and 3 provide equivalent information. Segments of type 2 are most informative since they represent a direct observation of the distance between a pair of chiasmata. The likelihood of the data $\mathcal{L}(y_i|\nu)$ is obtained by summing over the likelihood of all types of segments.

When $\nu = 1$, the waiting time follows an exponential distribution, which corresponds to the Poisson process and therefore no interference. Values of $\nu > 1$ tend create more even spacing of crossovers and correspond to positive interference.

### Gamma-sprinkling model

This model treats the position of crossovers as a mixture of two distinct processes – one, with interference ($\nu \neq 1$), with probability $1 - p$; and the other, without interference ($\nu \equiv 1$), with probability $p$. Although the motivation for this model is to account for the fact that some proportion of crossovers in many organisms arise via a non-interfering pathway, the model does *not* explicitly designate any specific crossover as interfering or non-interfering. The proportion of non-interfering crossovers ($p$) acts only as a weighting factor in the likelihood function. See Appendix A of Housworth and Stahl (2003) for details.

Simulations over a range of values for $(\nu, p)$ help give some intuition about the effect of each parameter on the distribution of inter-crossover distances. I focus here on chromosomes with at least two crossovers because it makes the effect most obvious. Each panel in **Figure B** shows the aforementioned distribution from simulation of 25 000 meiotic products of uniform length (100 cM). See that as $\nu$ increases, the spread of distribution of inter-crossover distances becomes smaller; that is, crossovers tend to be more evenly spaced. However, as $p$ increases, another mode appears in the lower tail of the distribution, corresponding to non-interfering crossovers that tend to appear closer to their neighbors. The theoretical distribution is *not* a simple mixture, but the contributions of the two pathways are visually clear.

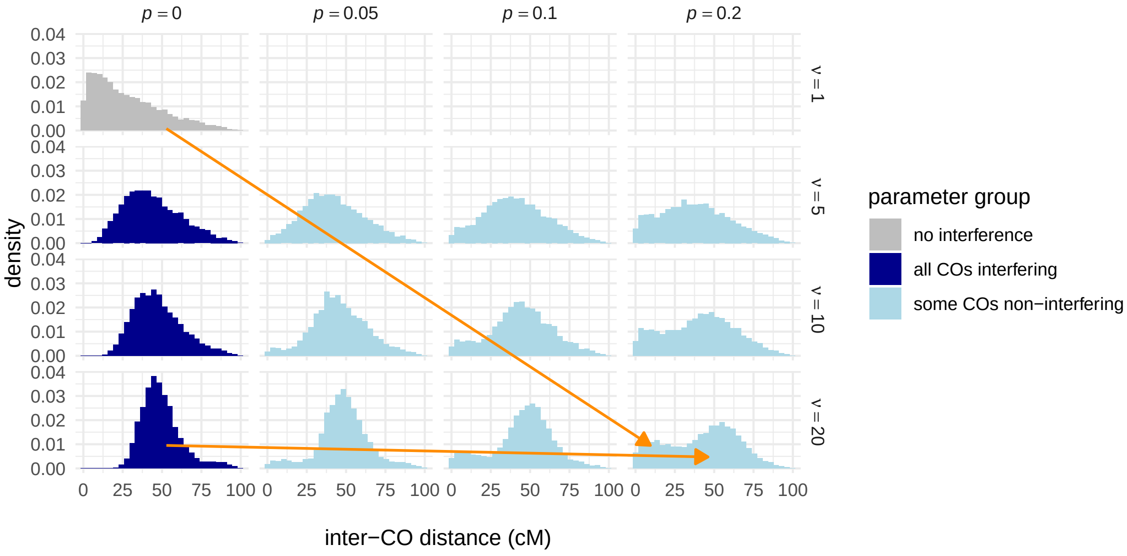


Figure B: **Simulated distribution of inter-crossover distance for varying $\nu$ and $p$.**
